# Stabilization of V1 interneuron-motor neuron connectivity ameliorates motor phenotype in a mouse model of ALS

**DOI:** 10.1101/2022.12.15.520568

**Authors:** Santiago Mora, Rasmus von Huth Friis, Anna Stuckert, Gith Noes-Holt, Roser Montañana-Rosell, Andreas Toft Sørensen, Raghavendra Selvan, Ilary Allodi

## Abstract

Loss of connectivity between spinal V1 inhibitory interneurons and motor neurons is found early in disease in the SOD1^G93A^ ALS mice. Such changes in premotor inputs can contribute to homeostatic imbalance of vulnerable motor neurons. Here, we show, for the first time, that stabilization of V1 synapses by overexpression of the Extended Synaptotagmin 1 presynaptic organizer increases motor neuron survival and ameliorates motor phenotypes, demonstrating that interneurons can be a potential target to attenuate ALS symptoms.

## Main

Somatic motor neurons are the ultimate output of the brain since they control movements by directly connecting to muscles. Their synchronized activation is regulated by a complex network of inhibitory and excitatory spinal interneurons ^1^. Hence, functional connectivity between motor neurons and their premotor circuits is a prerequisite for maintenance of inhibitory-excitatory balance and execution of movements. In the fatal disease Amyotrophic Lateral Sclerosis (ALS), somatic motor neurons degenerate, and subjects progressively lose the ability to perform movements. In our previous work ^2^, we showed that the spinal V1 inhibitory interneurons, positive for Engrailed-1 (En1) marker, lose their synapses onto the vulnerable fast-twitch fatigable motor neurons early in disease in the SOD1^G93A^ ALS mouse model ^3^. This preferential loss of inhibitory inputs onto fast-twitch fatigable motor neurons might contribute to their unbalanced excitability ^4^, leading to excitotoxicity and ultimately to their vulnerability to disease. Moreover, V1 inhibitory interneurons are known to control the speed of locomotion in vertebrates ^5^, and the loss of connectivity observed in the SOD1^G93A^ mice led to an onset of locomotor phenotype. Such phenotype is characterized by a reduction of speed and acceleration, a decrease in stride length and step frequency, and a hyperflexion of the hindlimbs ^2^. These symptoms were observed at a timepoint preceding motor neuron death and muscle denervation, and could be directly associated to loss of V1 inputs ^2, 5, 6^. Further evidence of alterations in inhibitory synapses has been reported also in a Fused in sarcoma (FUS) mouse showing Amyotrophic Lateral Sclerosis and Frontotemporal Dementia (FTD)-like phenotypes ^7–8^. Thus, inhibitory synaptopathy is not restricted to the SOD1^G93A^ mouse model. Interestingly, synaptic proteomics performed on postmortem tissue of C9ORF72 patients identified ~500 proteins with altered expression levels also within inhibitory synapses ^9^. Moreover, two recent studies showed that misprocessing of UNC13A mRNA strongly associates with ALS-FTD pathology caused by TDP43 downregulation ^10, 11^. The UNC13A gene plays a pivotal role in neurosecretion and is a fundamental component of neuron-to-neuron communication ^12, 13^, suggesting general synaptic dysregulations in the disease. Thus, the potential development of strategies directed to promote neurosecretion and synapse stabilization might be beneficial in the attempt to overcome such synaptopathy.

In the present study, we investigated if by stabilizing connectivity between spinal V1 inhibitory interneurons and motor neurons we could modify disease progression in the SOD1^G93A^ mice. To this aim, we overexpressed the human presynaptic protein Extended synaptotagmin 1 (Esyt1) specifically in V1 interneurons. Esyt1 was previously shown to promote synaptic growth and stabilization ^14^, and is preferentially expressed in neurons resistant to ALS ^15^. Moreover, Esyt1 transcript is downregulated in neurons within the ventral horn of the spinal cord of SOD1^G93A^ mice at postnatal day 63 (P63) (Figure 1a-d), the same timepoint at which we found decreased levels of the En1 transcript ^2^. To achieve V1 restricted overexpression, an adeno associated (AAV) serotype 8 virus was generated to overexpress Esyt1 upon *cre*-dependent recombination. The AAV8-hSyn-DIO-hEsyt1-W3SL virus was injected intra-spinally in SOD1^G93A^ mice crossed with En1^cre^ mice ^16^ (Figure 1e-f). Phenotype and genotype of the SOD1^G93A^;En1^cre^ mice, including copy number of the mutated gene, was evaluated and did not differ from mice expressing SOD1^G93A^ alone (Supplementary figure 1a-c). All four genotypes resulting from the crossing – WT, En1^cre^, SOD1^G93A^ and SOD1^G93A^;En1^cre^ – received bilateral injections in each of the L1-L3 lumbar segments (six in total) of 100 nl each at postnatal day 30. Virus was used at a final titer of 4×10^12^ vg/mL. L1-L3 segments were targeted since they are the first affected in SOD1^G93A^ mice ^17^, and responsible for the onset of locomotor phenotype ^2^. An AAV8-hSyn-DIO-mCherry-WPRE virus was also generated and injected as described above in the lumbar segment of the spinal cord of En1^cre^ and SOD1^G93A^;En1^cre^ mice to validate V1-restricted transduction (Figure 1e, g). Due to the large size of the Esyt1 insert (~3.3k bp), a fluorescent tag could not be added to the AAV8-hSyn-DIO-hEsyt1-W3SL and Esyt1 overexpression was analyzed utilizing an RNAscope probe recognizing the inverted vector sequence upon *cre-*recombination (Figure 1h). As expected, overexpression was specific to neurons within the ventral/medial areas of the spinal cord and restricted to *cre* mice (Figure 1h-j). Fluorescence was not detected in the WT injected mice (Figure 1k-m). Quantification of hEsyt1 revealed overexpression in 20% of En1^+^ neurons in the analyzed lumbar segments (Figure 1n).

**Figure 1.**
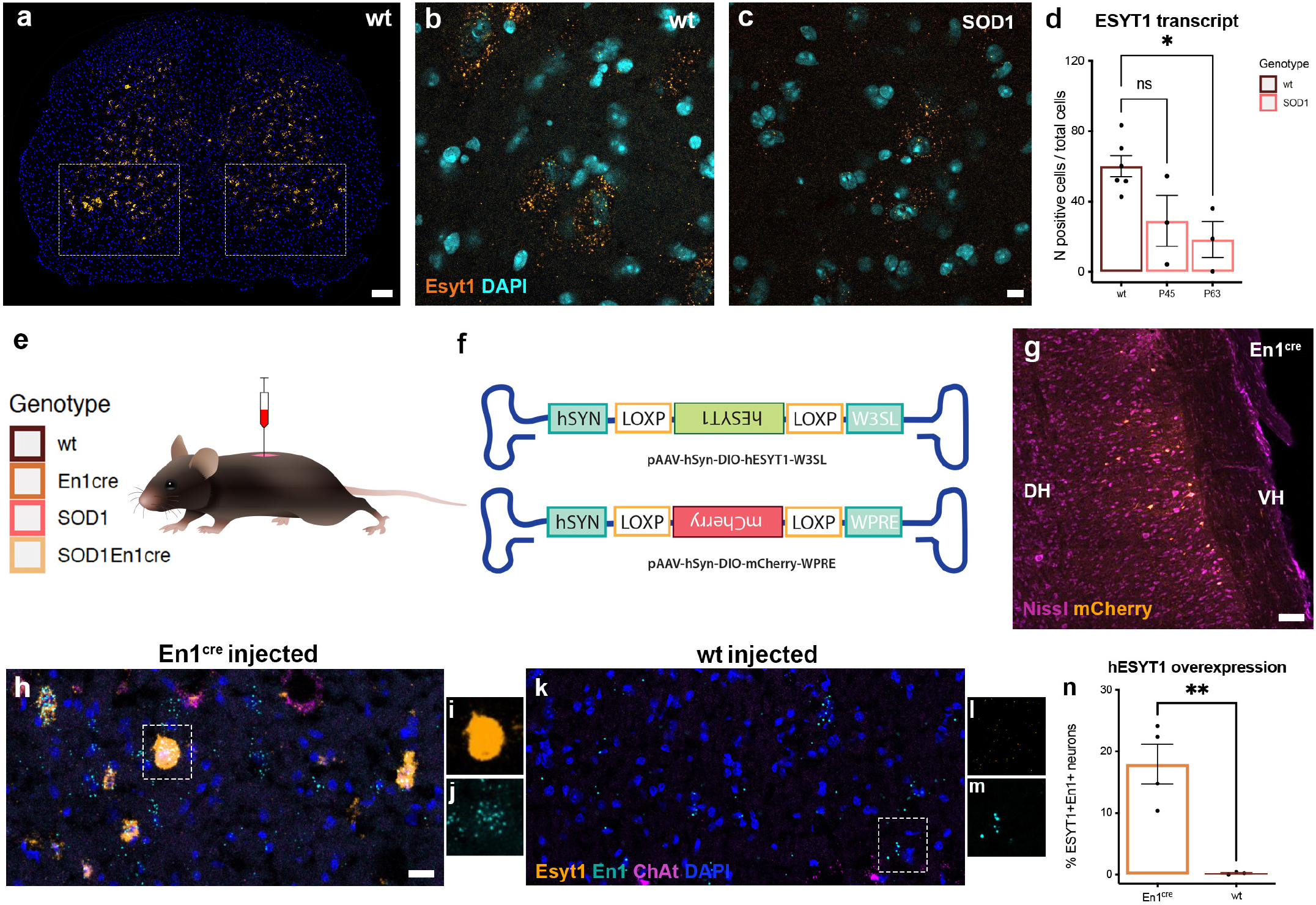
Esyt1 overexpression in SOD1^G93A^;En1^cre^ mice. a) Expression of Esyt1 transcript in the lumbar spinal cord of a WT mouse visualized by RNAscope. b) 40X magnification microphotograph showing Esyt1 expression in the ventral horn of the spinal cord in a WT mouse. c) Esyt1 is downregulated in neurons during early ALS progression in the ventral horn of the spinal cord. Esyt1 in orange, DAPI in light blue. d) Quantifications performed in WT and SOD1^G93A^ mice at postnatal day 45 and 63 show downregulation of Esyt1 transcript starting at P63 (Oneway ANOVA and Dunnett’s post hoc, P45 P=0.0714, P63 P=0.0191, N=3 per timepoint, per genotype). e) Cartoon depicting methodological approach in all experiments. Intraspinal injections delivering AAV vectors were performed in WT, En1^cre^, SOD1^G93A^ and SOD1^G93A^;En1^cre^ mice throughout the study, all genotypes were injected in L1-L3 spinal segments. f) Schematic of *cre*-dependent constructs designed to overexpress hEsyt1 and mCherry. g) Longitudinal section of the spinal cord of an En1^cre^ mouse upon overexpression of the AAV8-hSyn-DIO-mCherry virus validates successful *cre*-dependent expression in the ventral/medial areas of the cord. AAV8-hSyn-DIO-hEsyt1-W3SL driven overexpression was analyzed by RNAscope utilizing a probe recognizing the inverted viral construct upon *cre-*recombination. h) Expression of viral-hEsyt1 in En1^cre^ mice. i) Close-up microphotograph showing a neuron overexpressing hEsyt1. j) The same neuron is also positive for En1 transcript. k) hEsyt1 was not detected WT injected mice. l-m) En1 positive neuron negative for hEsyt1. n) Quantification of Esyt1 positive neurons in the lumbar spinal cord of En1^cre^ and WT mice, 20% of En1^+^ neurons are positive for the hEsyt1 (*t* test, P=0.0057, En1^cre^ N=4, WT N=3). Scale bar in a) = 100 μm, in b) = 50 μm and in h) = 100 μm. All graphs show mean values ± SEM.

Upon AAV8-hSyn-DIO-hEsyt1-W3SL administration in all four genotypes, changes in inhibitory synaptic density on motor neurons were investigated at postnatal day 112 (P112) (Figure 2a-e). This timepoint was chosen since significant motor neuron loss can be observed at P112 in the SOD1^G93A^ mice ^18^, and we previously showed a decrease in inhibitory synaptic inputs from P45 ^2^. Inhibitory synaptic inputs were visualized utilizing a VGAT antibody, while motor neurons were identified by ventral localization, size, and Chat staining (Figure 2a-d). VGAT synaptic density on motor neurons was reconstructed and corrected by motor neuron area (Figure 2a-d). Injected SOD1^G93A^;En1^cre^ mice exhibited significant increase in synaptic density when compared to injected SOD1^G93A^, similar to injected WT and En1^cre^ conditions (Figure 2e). Motor neurons were also quantified at the same timepoint in all four AAV8-hSyn-DIO-hEsyt1-W3SL injected genotypes (Figure 2f-l). Here, only neurons over 28 μm in diameter within the ventral horn of the spinal cord (putative of fast motor neurons) were quantified. SOD1^G93A^; En1^cre^ mice showed an increased number of spared motor neurons compared to SOD1^G93A^ mice after AAV8-hEsyt1 overexpression. A trend in reduction of larger motor neurons in En1^cre^ control mice upon hEsyt1 overexpression was observed. Analysis of motor neuron areas of control En1^cre^ mice with and without hEsyt1 overexpression demonstrated a shrinkage of motor neurons at P112 (Figure 2m).

**Figure 2.**
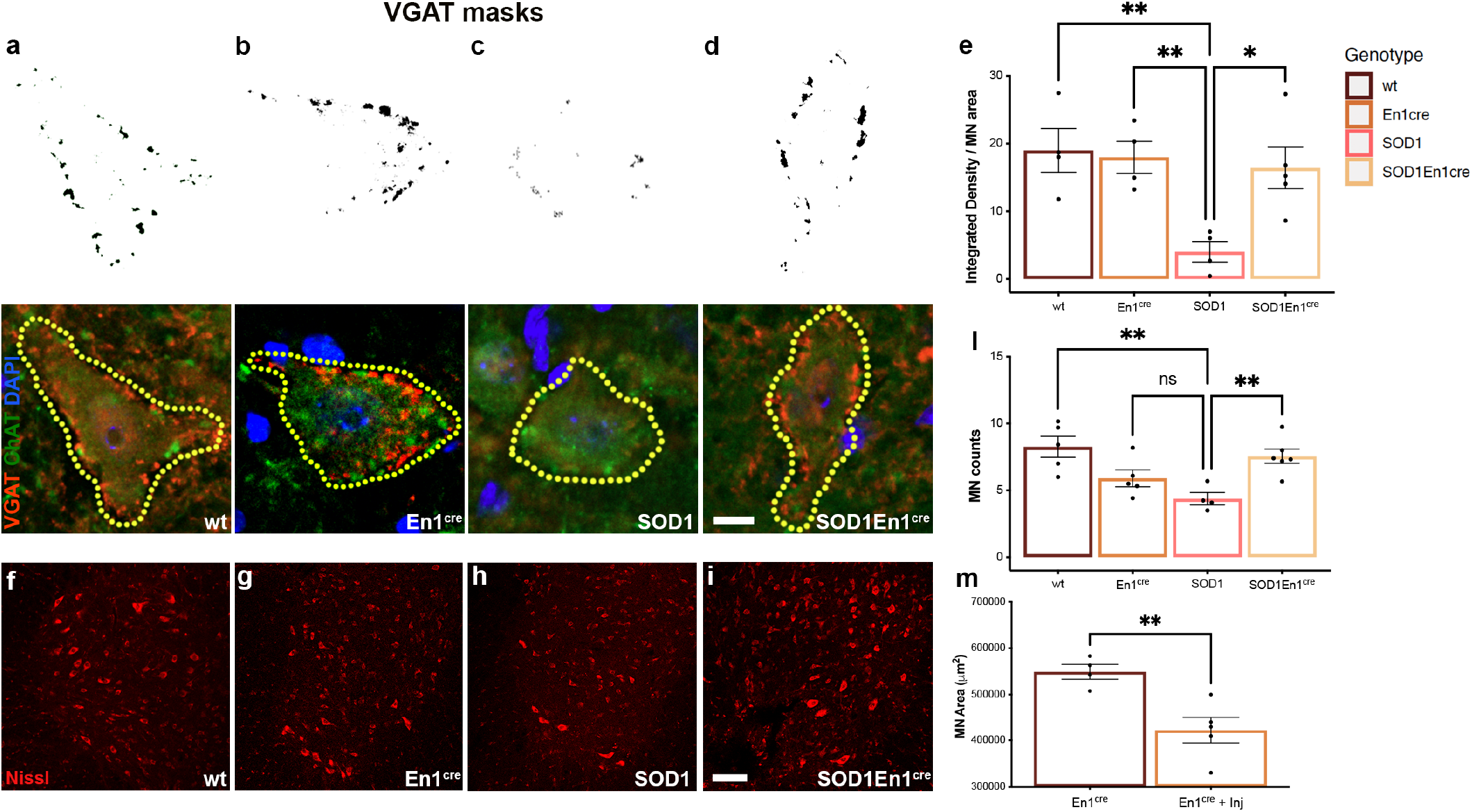
Esyt1 overexpression increases motor neuron survival and synaptic density onto spared motor neurons. Quantifications of inhibitory synapses on spared motor neurons at postnatal day 112 upon AAV8-hSyn-DIO-hEsyt1 injections. Synaptic densities are normalized by motor neuron area. Masks in a) and b) show synaptic density in WT and En1^cre^ mice respectively, c) and d) show differences in synaptic densities between SOD1^G93A^ and SOD1^G93A^;En1^cre^ mice. Microphotographs show examples of quantified motor neurons in the different genotypes. VGAT in red, ChAT in green, DAPI in blue. Scale bar in g) = 50 μm. e) Synaptic density in SOD1^G93A^;En1^cre^ is significantly increased compared to SOD1^G93A^ and comparable to the synaptic densities of WT and En1^cre^ mice (One-way ANOVA and Dunnett’s post hoc WT P = 0.0060, En1^cre^ P = 0.0097, SOD1 ^G93A^;En1^cre^ P = 0.0149, N = 4). Motor neuron quantifications performed at postnatal day 112 in f) WT, g) En1^cre^, h) SOD1^G93A^ and i) SOD1^G93A^;En1^cre^ mice. Fluoro-Nissl in red. Scale bar in i) = 50 μm. l) SOD1^G93A^;En1^cre^ mice show increased motor neuron survival upon Esyt1 overexpression when compared to the SOD1^G93A^ mice (One-way ANOVA and Dunnett’s post hoc SOD1^G93A^; En1^cre^ P = 0.0080, WT P = 0.0023, N = 5). A minimum of 170 motor neurons per condition was quantified. En1^cre^ mice overexpressing hEsyt1 show a trend in lower number of large motor neurons in the lumbar spinal cord (One-way ANOVA and Dunnett’s post hoc, En1^cre^ P = 0.2732, N = 5). m) Comparison of motor neuron size in En1^cre^ non-injected and En1^cre^ injected mice with AAV8-hSyn-DIO-hEsyt1 shows shrinkage of motor neurons upon hEsyt1 overexpression *(t* test, P = 0.0073, En1^cre^ non-injected N = 4, En1^cre^ injected N = 5). All graphs show mean values ± SEM.

Finally, we analyzed motor phenotypes after hEsyt1 overexpression in all four genotypes by placing the mice on a treadmill at a speed of 20 cm/s, equivalent to a fast walk ^19^. Videos were recorded from ventral and lateral views (Figure 3a;j). Our previously published data showed that ~40% of SOD1^G93A^ mice cannot cope with such speed by P63. Hence, we investigated if the increased synaptic connectivity upon hEsyt1 overexpression could ameliorate SOD1^G93A^ motor phenotype. Following a brief training period, mice were assessed once a week from postnatal day P49 until P112. Videos were analyzed using the DeepLabCut marker-less pose estimation tool ^20^ as previously described ^2^. SOD1^G93A^ mice showed a significant reduction in locomotor performance and, consistently with our previous data, 37.5% showed an onset of locomo-tor phenotype by P63 (Supplementary Figure 2e). However, upon hEsyt1 overexpression only 16.7% of the SOD1^G93A^; En1^cre^ mice had an onset of locomotor phenotype by P63, and 50% of them could perform the task by P112 (SOD1^G93A^ median = P70, SOD1^G93A^;En1^cre^ median = P105). Here, average speed, step frequency, stride length and peak acceleration were analyzed longitudinally (Figure 3b-e). SOD1^G93A^;En-1^cre^ mice showed an amelioration in average speed from P63 (Figure 3b), step frequency from P70 (Figure 3c), and stride length from P84 (Figure 3d). However, we did not observe changes in peak acceleration (Figure 3e). When assessing the final timepoint (P112) SOD1^G93A^;En1^cre^ mice generally outperformed SOD1^G93A^ mice, and half of them showed preserved average speed (Figure 3f). Amelioration was also observed for step frequency (Figure 3g), and stride length (Figure 3h), while peak acceleration remained unchanged (Figure 3i). Intralimb kinematics was performed by analyzing changes in degree amplitudes of four angle joints: hip, knee, ankle, and foot (Supplementary Figure 3 and Supplementary video 1 and 2). 15 individual steps per trial were analyzed and the mean amplitude for each mouse was calculated. Hindlimb hyperflexion, a key element of the phenotype resulting from the loss of V1 synaptic inputs (Supplementary Figure 3), was improved in the SOD1^G93A^;En1^cre^ mice upon hEsyt1 overexpression for the foot and the ankle angles, but not for the hip and the knee (Figure 3k-n). Moreover, comparisons of lateral view recordings between SOD1^G93A^ and SOD1^G93A^;En1^cre^ mice revealed that the latter could better support their body weight as they maintained their bodies at a greater distance from the belt of the treadmill (Supplementary video 1 - kinematics of SOD1^G93A^ and Supplementary video 2 – kinematics of SOD-1^G93A^;En1^cre^).

**Figure 3.**
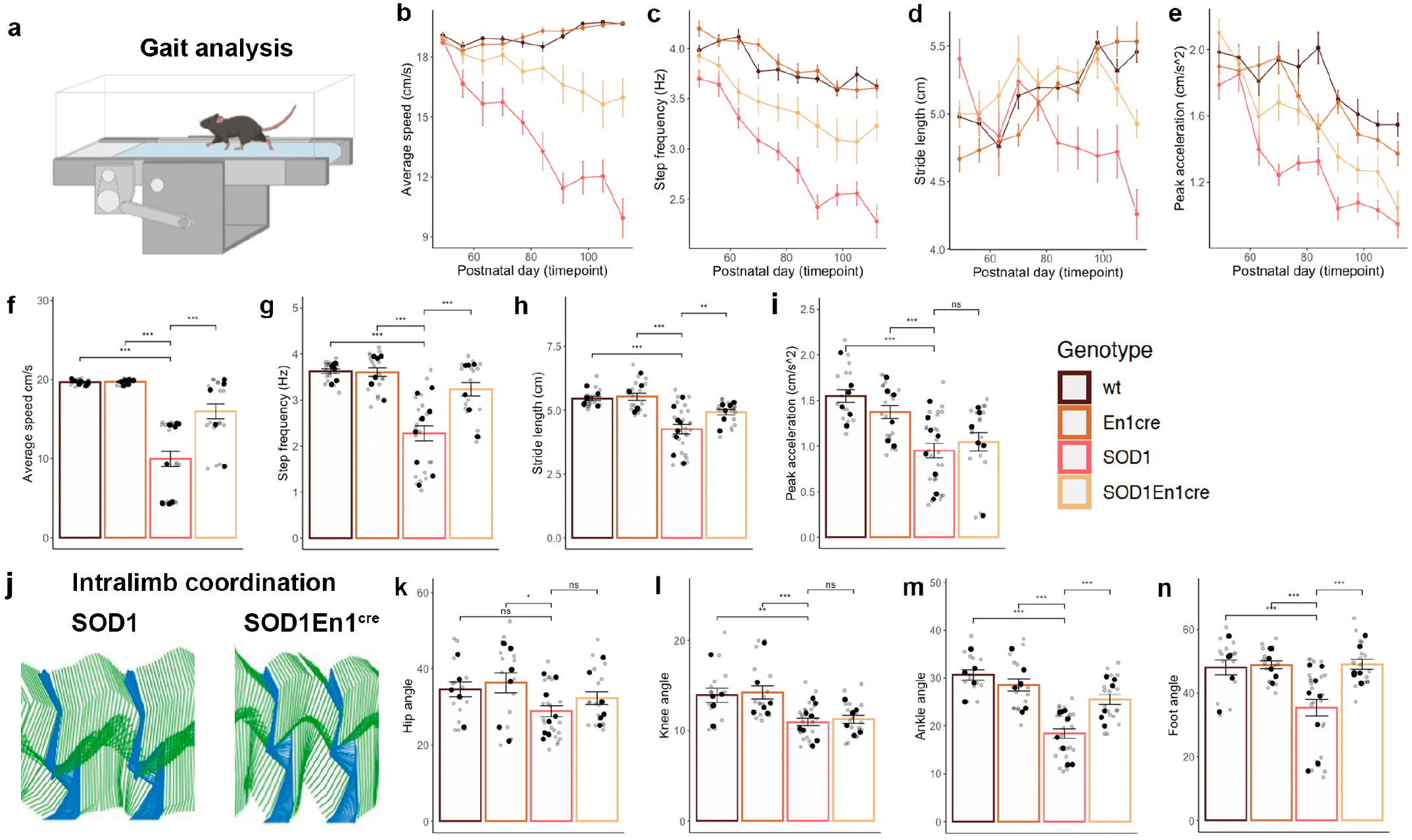
Amelioration of locomotor phenotype in SOD1^G93A^ mice upon hEyt1 overexpression. a) Cartoon depicting treadmill paradigm. b) Average speed analyzed between day 49 and 112 shows amelioration in SOD1 ^G93A^;En1^cre^ mice from P63 (Two-way ANOVA and Dunnett’s post hoc, P = 0.0257), c) step frequency from P70 (Two-way ANOVA and Dunnett’s post hoc, P = 0.0318), and d) stride length from P84 (Twoway ANOVA and Dunnett’s post hoc, P = 0.0096). e) Peak acceleration does not change upon Esyt1 overexpression (Two-way ANOVA and Dunnett’s post hoc, N = 6-8 mice per condition; all quantifications were performed in triplicates). At P112 timepoint f) average speed is higher in SOD1^G93A^;En1^cre^ compared to SOD1^G93A^ mice (One-way ANOVA and Dunnett’s post hoc, SOD1^G93A^;En1^cre^ vs SOD1^G93A^ P = 1.8e-07), as well as g) step frequency (One-way ANOVA and Dunnett’s post hoc, P = 3.1e-06) and h) stride length (One-way ANOVA and Dunnett’s post hoc, P = 0.0032), however i) peak acceleration remains unchanged (One-way ANOVA and Dunnett’s post hoc, P = 0.7406) (WT N = 6; SOD1^G93A^ N = 8, En1^cre^ N = 6, SOD1^G93A^;En1^cre^ N = 6, all quantifications were performed in triplicates). j) Stick figures depict intralimb coordination in SOD-1^G93A^ and SOD1^G93A^;En1^cre^ mice. Two full cycles are visualized for SOD1^G93A^and SOD1^G93A^;En1^cre^ mice upon AAV8-hSyn-DIO-hEsyt1 injections. Stance phase in blue and swing phase in green. Changes in joint angles were analyzed for k) hip angle, l) knee angle, m) ankle angle and n) foot angle. Significant changes were found for the foot angle (One-way ANOVA and Dunnett’s post hoc, foot angle P = 4.1e-05) and the ankle angle (One-way ANOVA and Dunnett’s post hoc, foot angle P = 2.8e-05) upon hEsyt1 overexpression (WT = 5, SOD1^G93A^ N = 8, En1^cre^ N = 6, SOD1^G93A^;En1^cre^ N = 6, all quantifications were performed in triplicates). All graphs show mean values ± SEM, averages values in f-I and k-n are shown in black, technical triplicates are shown in gray.

Altogether, these results indicate that motor impairment in SOD1^G93A^ mice can be attenuated by stabilization of synaptic inputs between V1 interneurons and motor neurons. By overexpression of Esyt1 presynaptic protein in V1 interneurons, we were able to increase motor neuron survival and to slow down disease progression. It has been previously hypothesized that therapies targeting specific microcircuit dysfunctions might help slowing down the course of the disease ^21^. The present work supports this hypothesis and indicates that the V1 interneurons-motor neuron circuit can be a potential target for treatment in ALS. Electrophysiological studies investigating recurrent inhibition support general inhibition dysregulation in ALS patients, and reduced inhibition in patients showing initial motor weakness in the lower limbs ^22^. Thus, further effort will be required to identify a potential therapeutic window for targeting premotor circuits at the appropriate timepoint to slow down the disease. Moreover, while specific intraspinal delivery within the lumbar segments of the spinal cord was found beneficial in this study, it remains to be inves-tigated if systemic overexpression of hEsyt1 in En1 positive cells could further improve disease phenotype and survival. The surprising maladaptive changes observed in healthy En-1^cre^ mice upon Esyt1 overexpression, leading to motor neuron shrinkage, suggest the need for targeted administration. En1 is expressed in neuronal populations outside the spinal cord (e.g., dopaminergic neurons) that could be affected by hEs-yt1 overexpression. Hence, the use of specific enhancers ^23^ or synthetic synaptic organizers ^24^ could be viable alternatives for selective targeting of affected circuits.

## Methods

### Ethical Permits and Mouse strains

All experiments were in accordance with the EU Directive 20110/63/EU and approved by the Danish Animal Inspectorate (Ethical permits: 2018-15-0201-01426 and 2022-15-0201-01164). SOD1^G93A^ (B6.Cg-Tg(SOD1-G93A)1Gur/J) stock no: #004435 were retrieved from Jackson Laboratory, while En1^cre^ mice, kept on a C57BL6/J background, were provided by Assistant Prof. Jay Bikoff (St. Jude Children’s hospital, St Louis, Texas USA). Mice were genotyped using DNA extracted from earclipping and the following primers were used: for the SOD1^G93A^ gene, 5’-CAT CAG CCC TAA TCC ATC TGA-3’ and 5’ -CGC GAC TAA CAA TCA AAG TGA-3’, while for the En1^cre^ gene, 5’-GAG ATT TGC TCC ACC AGA GC-3’ and 5’-AGG CAA ATT TTG GTG TAC GG-3’. Copy number of the mutated SOD1 gene was quantified by qPCR utilizing the primers 5’-GGG AAG CTG TTG TCC CAA G-3’ and 5’-CAA GGG GAG GTA AAA GAG AGC-3’ for the SOD-1^G93A^ gene. The qPCR reaction was performed as suggested by the mice supplier. All mouse strains were bred with congenic C57BL6/J mice, stock no: #000664 (Jackson Laboratory). Mice were housed according to standard conditions with *ad libitum* feeding, constant access to water, and a 12:12 hour light/dark cycle. All mice, including multiple crossing, were genotyped, tested for copy number, and phenotype was assessed, including weekly weight. For survival experiments, the humane endpoint was defined as a weight loss of 15% and/ or functional paralysis in both hind limbs together with inability to perform a righting test <20 seconds. Mice of both genders were included in the study and non-transgenic SOD1^G93A^ littermates were used as controls.

### Adeno-associated Viral Vector production

The AAV overexpressing hEsyt1 was developed utilizing a pAAV-hSyn-DIO[hCAR]off-[hM4Di-mCherry]on-W3SL backbone construct containing the W3SL cassette (Addgene plasmid #111397). First, hCAR and hM4di-mCherry were exerted by double digestion utilizing *Asc1*/*Nhe1* restriction enzymes, the mCherry fluorescent tag was exerted, and Kozak sequences inserted by PCR. cDNA of the human Extended Synaptotagmin 1 (hESyt1) was purchased by Dharmacon (#MHS6278-202826307) and insert amplified by PCR utilizing Primers 5’-TAG CAG GCG CGC CCT AGG AGC TGC CCT TGT CC-3’ and 5’-GAG TCT CTA GAG CCA CCA TGG AGC GAT CTC CAG GAG AG-3’ containing a Kozak sequences at the start codon position. The PCR product was double digested utilizing *XbaI*/*AscI* restriction enzymes and ligated into the pAAV backbone. Both backbone construct and insert were sequenced to validate successful sequence-content and orientation. The pAAV-vector, encoding for hEsyt1, the pHelper carrying the adenovirus-derived genes, and pAAV-Rep-Cap (carrying the AAV2 replication and AAV8 capsid genes) were co-transfected in 293-cells with Dulbecco’s Modified Eagle Medium (DMEM) supplemented with 10% fetal bovine serum (FBS) and 1% penicillin-streptomycin (all from GeneMedi). The following day, medium was replaced. Medium containing viral particles was harvested approximately 72 h later through low-speed centrifugation at 1.500 g in an Eppendorf centrifuge (Eppendorf 5810R, Hamburg, Germany) for 5 min. Cell pellet was resuspended in 10 mM tris(hydroxymethyl)aminomethane hydrochloride (pH 8,5) lysis buffer with subsequent freeze/thaw cycles. Supernatant containing the harvested AAV-particles was collected after 10 min centrifugation at 3.000 g and filtered through a 0,22 μm filter. Concentration of the filtered supernatant occurred by ultrafiltration and several centrifugation steps. The pellet containing the viral particle was resuspended in 0,1 M PBS pH 7,4 and aliquoted and stored at −80°C until further use. The AAV8-hSYN-DIO-mCherry vector was purchased from Addgene (plasmid #50459: Bryan Roth lab). The two viral vectors had equal titers of 4.0×10^12^ vg/ml.

### Intraspinal injections and viral delivery

To achieve *cre*-dependent overexpression of hEsyt1 in V1 interneurons, En1^cre^ mice were used and crossed with SOD-1^G93A^. For anatomical and behavioral experiments performed in this study, all littermates (including WT, SOD1^G93A^;En1cre, En1^cre^, and SOD1^G93A^ mice) were injected with AAV8-hSyn-DIO-hEsyt1-W3SL at postnatal day 30 by an experimenter blind to the genotype. The four injected genotypes were included in the study, to exclude off target effects of the viral vector and changes due to the surgical procedure. For intra-spinal injections mice were anaesthetized with 2% isoflurane and the lumbar level of the spinal cord was exposed for stereotaxic injections (Neurostar). A small incision was performed with micro-scissors between the vertebrae to deliver the virus in the L1, L2 and L3 spinal segments. For visualization, virus was mixed with 4% fast green (Invitrogen) dissolved in saline and injected using a glass micropipette at a rate of 100 nl/min. The micropipette was kept in place for 2 min after viral delivery to avoid backflow. Bilateral injections of 100 **μ**l AAV8-hSyn-DIO-hEsyt1-W3SL virus were performed. Additionally, similar injections of 100 **μ**l of a control AAV8-hSyn-DIO-mCherry-WPRE were conducted to validate successful *cre*-recombinase upon injection. Pre-operatively mice were treated with a subcutaneous injection of buprenorphine diluted in saline at a concentration of 0.3 mg/ml. Moreover, post-operatively pain relief was administered for three days using a similar dose of buprenorphine (Temgesic) mixed in DietGel Boost (ClearH_2_O).

### RNAscope protocol

For RNAscope assay, mice were anaesthetized with an overdose of Pentobarbital (250 mg/kg) and sacrificed by decapitation. Spinal cords were dissected, snap frozen in isopentane (2-methylbutane, Uvasol, Merck) and kept in dry ice for cryoprotection. Then, spinal cords were coronally or longitudinally sectioned at 12 μm-thickness by using a cryostat (Thermo Fisher Scientific), collected on Superfrost Plus slides (Thermo Fisher Scientific) and stored at −80°C until further processing. Samples were pre-treated and processed using the RNAscope Multiplex Fluorescent v2 Assay following the supplier’s protocol (Advanced Cell Diagnostics - ACD). Briefly, sections were fixed in 4% PFA (HistoLab) for 15 minutes at room temperature, washed in DPBS and dehydrated with sequential ethanol steps. Then, they were incubated with RNAscope hydrogen peroxide solution for 10 minutes at room temperature, rinsed in DEPC-treated water, treated with RNAscope protease IV for 30 minutes at room temperature, and washed in PBS before in situ hybridization. Probes were hybridized for 2 hours at 40°C in HybEZ oven (ACD), samples were stored in 5X Saline Sodium Citrate (SSC) overnight at room temperature, followed by incubation with signal amplification and developing reagents according to the manufacturer’s instructions. Probes were purchased from ACD: Hs-ESYT1-C1 (catalog #540391), pAAV-hESYT1-WPRE-O1-C3 customized probe (catalog #1062751-C3), Mm-En1-C1 (catalog #442651), Mm-Chat-C2 (catalog #408731-C2). The hybridized probe signal was visualized and captured on a Zeiss LSM 900 confocal microscope with a Plan-Apochromat 40x oil objective – NA = 1,4 or 20x air objective – NA = 0,8, zoom = 1. Image analysis of RNAscope microphotographs was performed on tiled images of the ventral region of each section. A total of 10 tiled images were analysed per mouse and condition. Quantification of Esyt1 positive cells was performed in an automated manner using the open-sourced bio-image analysis software QuPath (version 0.2.3) ^25^. For cell segmentation and quantification of positive cells with the first probe, the “Positive cell detection” analysis function was used – settings shown in Supplementary Table 1. For the second probe, a triple filtering process was performed: first, with the *Create single measurement classifier* tool, a single filter was created for each marker, and the output was defined depending on the cells falling above or below the threshold (for ChAT: “Ignore” and “Other”; for En1 and ESYT1: “Positive” and “Negative”, respectively). Then, with the *Create composite classifier* tool, the number of En1+ cells expressing or not ESYT1 was calculated. Quantifications are presented as total amount of positive cells/total cells (Hs-ESYT1-C1 probe) or total amount of double positive cells/total amount of En1 positive cells (pAAV-hESYT1-WPRE-O1-C3 probe).

### Immunohistochemistry

Mice were injected with Pentobarbital (250 mg/kg) and transcardially perfused with pre-chilled phosphate buffered saline (PBS, Gibco) followed by pre-chilled 4% paraformaldehyde (PFA, HistoLab). Spinal cord was dissected, postfixed for 60 min in cold 4% PFA and cryoprotected in PBS 30% sucrose for 48 hours at 4°C. For immunofluorescence coronal and longitudinal sections of the lumbar spinal cord were sectioned at 30 **μm-thickness. Sections were** collected on Superfrost Plus slides (Thermo Fisher Scientific) and stored at −20°C for further processing. Slides were then washed for 10 minutes in PBS at room temperature and blocked for 1 hour in PBS with 0.1% Triton-X100 (PBS-T, Sigma Aldrich) and 1.5% donkey serum (Invitrogen). Sections were incubated for 24-48 hours at 4 °C in primary antibodies diluted in blocking solution. The following antibodies were used: DS Red (1:1000, Rabbit, Takara-Clontech), ChAT (1:300, Goat, Millipore), VGAT (1:1000, Rabbit, Millipore). Slides were washed three times in PBS at room temperature for 10 minutes each and then incubated for 1 hour with appropriate secondary antibodies (1:500, Alexa Fluor 488, 568, Invitrogen) diluted in blocking solution. Subsequently, another three 10-minute PBS washes, and counterstaining was performed with NeuroTrace 640 (1:200, Invitrogen) or Hoechst (1:2000, Invitrogen). After two 5-minute washes is PBS, slides were dried, and cover slipped using ProLong Diamond Antifade Mountant (Invitrogen). Microphotographs were obtained utilizing either Zeiss LSM 700 or 900 confocal microscopes, using a Plan-Apochromat 20X objective – NA = 0,8, zoom = 0,5. For motor neuron survival and synaptic density quantifications, one image of the ventral region of each hemisection was acquired. Quantifications were conducted utilizing Fiji software. Larger motor neurons were detected and quantified based on localization, shape, and cell diameter of 28 μm. For synaptic density quantifications, microphotographs acquired with a 20x objective (zoom = 1), were transformed to a grayscale 8-bit images and quantified after applying a threshold for signal intensity/background correction. Images with a higher background/noise ratio were excluded. ROI were drawn around the motor neurons of interest that showed clear soma staining and visible nucleus. Pixel contained in the area of interest were quantified with the Fiji function Analyze particles and masks of all quantified images were created to validate the analysis. Integrated intensity of pixels obtained with Fiji was normalized by the motor neuron area.

### DigiGait Treadmill Test

Locomotor performance was assessed using the DigiGait motorized transparent treadmill (Mouse Specifics, Inc.), which allows recording of mice from a ventral and lateral view. Mice were trained one week prior, then tested weekly from P49 until P112 on the treadmill at a speed of 20 cm/s for analysis of disease progression. After a 2-minute acclimatization to the treadmill, mice were recorded at a speed of 20 cm/s for 10 seconds in 3 consecutive trials with 2-minute rest periods between recordings. Belt speed was adjusted to 15, 10 or 5 cm/s as needed, depending on the locomotor capability.

### Kinematic Analysis

The videos captured during the treadmill experiments were analysed utilizing DeepLabCut ^20^ (DLC) software and an optimized model of the one previously described in Allodi et al. 2021. Briefly, eleven digital markers were placed on the mice and the tracks obtained by DLC were further analysed to extract the locomotor measures reported in the article. The reported measures are speed, acceleration (described as sudden increase or decrease in the speed), coordination (used to estimate the *stride length* and *step frequency).* The lateral view videos simultaneously captured during the treadmill experiments were analysed with a second DeepLabCut model. Six image markers (corresponding to iliac crest, hip, knee, ankle, foot, and toe) on the left hindlimb were tracked using this model. The tracked positions of the markers from each frame were further analysed to obtain four joint angles (at the hip, knee, ankle and foot) within each step cycle, as shown in Figure 3 (panels – k,l,m,n).

### Statistics

Esyt1 quantification, hEsyt1 overexpression, motor neuron quantification (MNq), synaptic density data, and MN size were analyzed by a One-way Analysis of variance (ANOVA) with number of positive cells/total cells (Esyt1 quantification), number of double positive cells/total amount of En1 positive cells (hEsyt1 overexpression), number of positive cells (MNq), and Integrated density/microns (Synaptic density) and motor neuron size (MN size), as dependent variable, and Strain (wt, SOD1^G93A^, SOD1^G93A^; En1^cre^, En1) (hEsyt1 overexpression, MNq and SD) or condition (En1^cre^/Inj; wt/ Inj) (Esyt1 quantification) (En1^cre^; En1^cre^/Inj) (MN size) as between-groups factor. When appropriate, post hoc comparisons with Dunnett’s test correction were performed. Moreover, statistical significance was set at P < 0.05, and effect size is reported when appropriate in figure legends: Partial eta-squared values are reported and considered as small (0.01), medium (0.06), or large (0.14). All analyses were performed using JASP©software (JASP Team, University of Amsterdam, version 0.16). For comparison of locomotor phenotype onset-curves, Log-rank (Mantel-Cox) test was employed. One-way ANOVA with subsequent post-hoc Dunnet’s test were used to compare differences between genotypes at single timepoints. The genotypes were compared against SOD1 as the reference group, to compare intervention and controls against the diseased mice. When considering multiple variables (genotype and time), two-way ANOVA was used. Twoway ANOVA tests were followed by Dunnett’s test post-hoc. The same approach was used for both ventral and lateral video data. All results are expressed as mean ± SEM, reported n values represent distinct biological replicates. P < 0.05 was considered statistically significant. Asterisks in figures and figure legends are * P < 0.05, ** P < 0.01 or *** P < 0.001.

Statistical analysis of behavior was performed in R Studio utilizing packages pacman, tidyverse, DescTools, ggsignif, emmeans, multcomp, and broom. No statistical methods were used to pre-determine sample sizes; our sample sizes are similar to those reported in previous publications ^2^. Data collection was performed blind to the conditions of the experiments although, in behavioural assessment, some ALS mice could be recognized at later stages. One mouse was excluded from the study since showed aberrant locomotor phenotype upon intraspinal injection, all other injected mice were included in the study.

## Supporting information

Kinematic analysis of a SOD1G93A;En1cre mouse at P112 at a speed of 15 cm/s (average phenotype)

Kinematic analysis of a SOD1G93A mouse at P112 at a speed of 10 cm/s (average phenotype)

## Author contribution

Conceptualization I.A., Methodology I.A., S.M., R.F., A.S., R.M.R, R.S., Viral vector production A.T.S, G.N.H., I.A., R.F. Data Analysis S.M., R.F., A.S., Supervision and Funding acquisition I.A.

## Acknowledgements

We thank Prof. Ole Kiehn, Department of Neuroscience – University of Copenhagen, for the access to the surgery room and the use of the DigiGait treadmill. We acknowledge the Core Facility for Integrated Microscopy, at the Faculty of Health and Medical Science of University of Copenhagen and the Department of Experimental Medicine, especially Dr. Pablo Hernandez-Varas (CFIM) and Alex Soelberg Laugesen (AEM). This work was supported by the Lundbeck Foundation (I.A.), the Louis-Hansen Foundation (I.A. and R.M.R.), the Danish Society for ALS (I.A.), the Laege Sofus Carl Emil Friis og hustru Olga Doris Friis’ foundation (I.A.) and the Danish Society for Neuroscience (R.F. and A.S.).

## Data availability

The data that support the findings of this study are available from the corresponding authors upon reasonable request.

## Code availability

The code used to analyze data, produce figure content and videos is available at: https://github.com/Allodi-Lab/hEs-yt1_spinal_injection_gait_analysis

## Supplementary information

**Supplementary figure 1.**
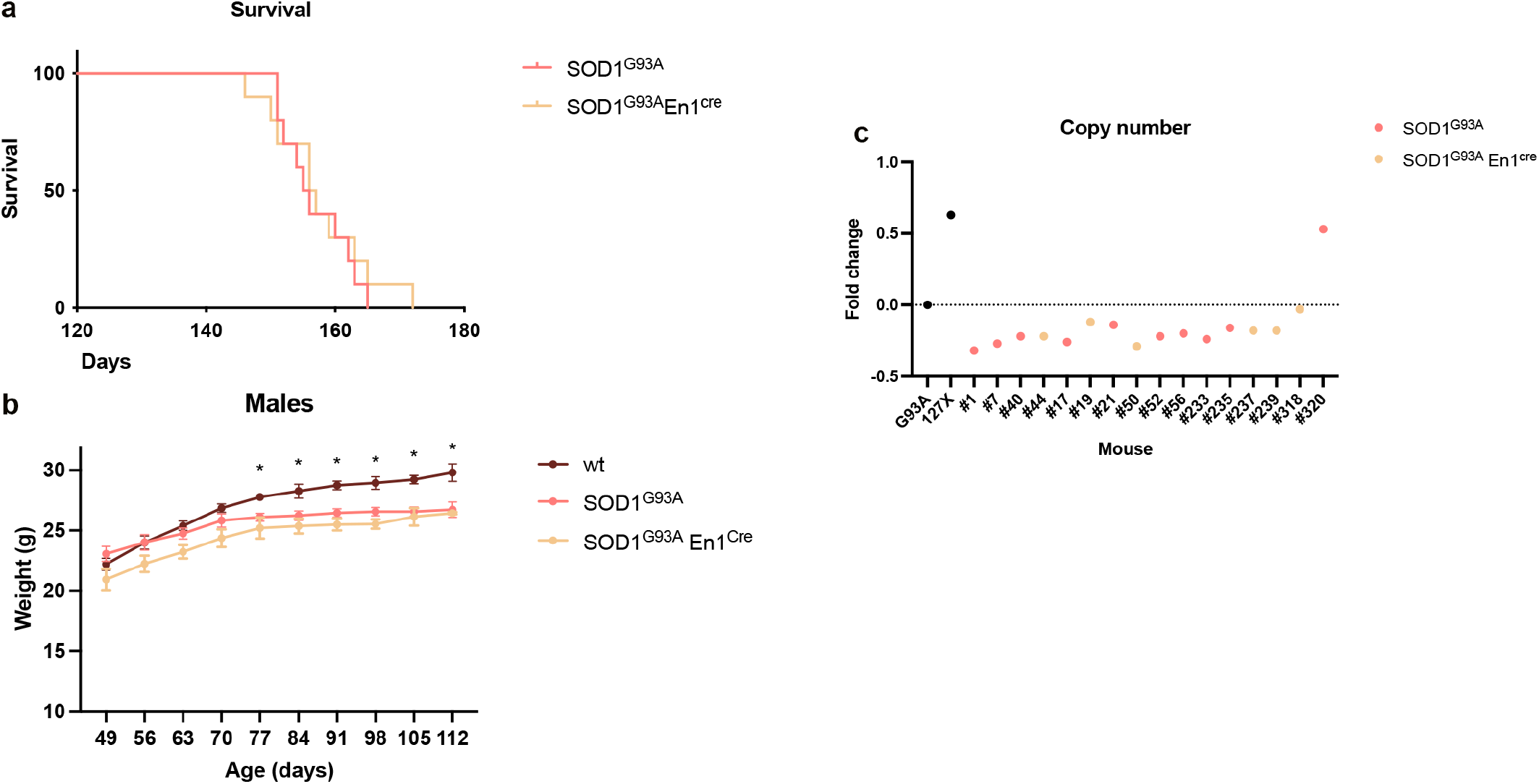
a) Kaplan Meyer survival curve comparing SOD1^G93A^;En1^cre^ to SOD1^G93A^ survival upon crossing. No significant differences were observed between the two strains (Log-rank (Mantel-Cox) test, P=0.5926, df=1, Chi square=0.2863; SOD1^G93A^;En1^cre^ N=10 mice, SOD1^G93A^ N=10 mice). b) Weights of SOD1^G93A^ and SOD1^G93A^;En1^cre^ males compared to WT mice. After crossing, SOD1^G93A^;En1^cre^ do not differ from SOD1^G93A^ mice but both strains decrease in weight compared to WT littermates (two-way ANOVA and Dunnett’s post hoc, F(18,110)=1.704, P=0.0489, P77 SOD1^G93A^ P=0.0307, SOD1^G93A^;En1^cre^ P=0.0046, WT N=5 mice, SOD1^G93A^ N=6 mice, SOD1^G93A^;En1^cre^ N=3 mice). c) Copy number of human SOD1 mutations carried by experimental mice. SOD1^G93A^ fold change in pink, SOD1^G93A^;En1^cre^ in beige. First black dot depicts positive control - SOD1^G93A^ founder carrying 25 copies of the mutated gene, second black dot depicts negative control - SOD1^127X^ carrying 19 copies of the mutated gene. All graphs show mean values ± SEM.

**Supplementary figure 2.**
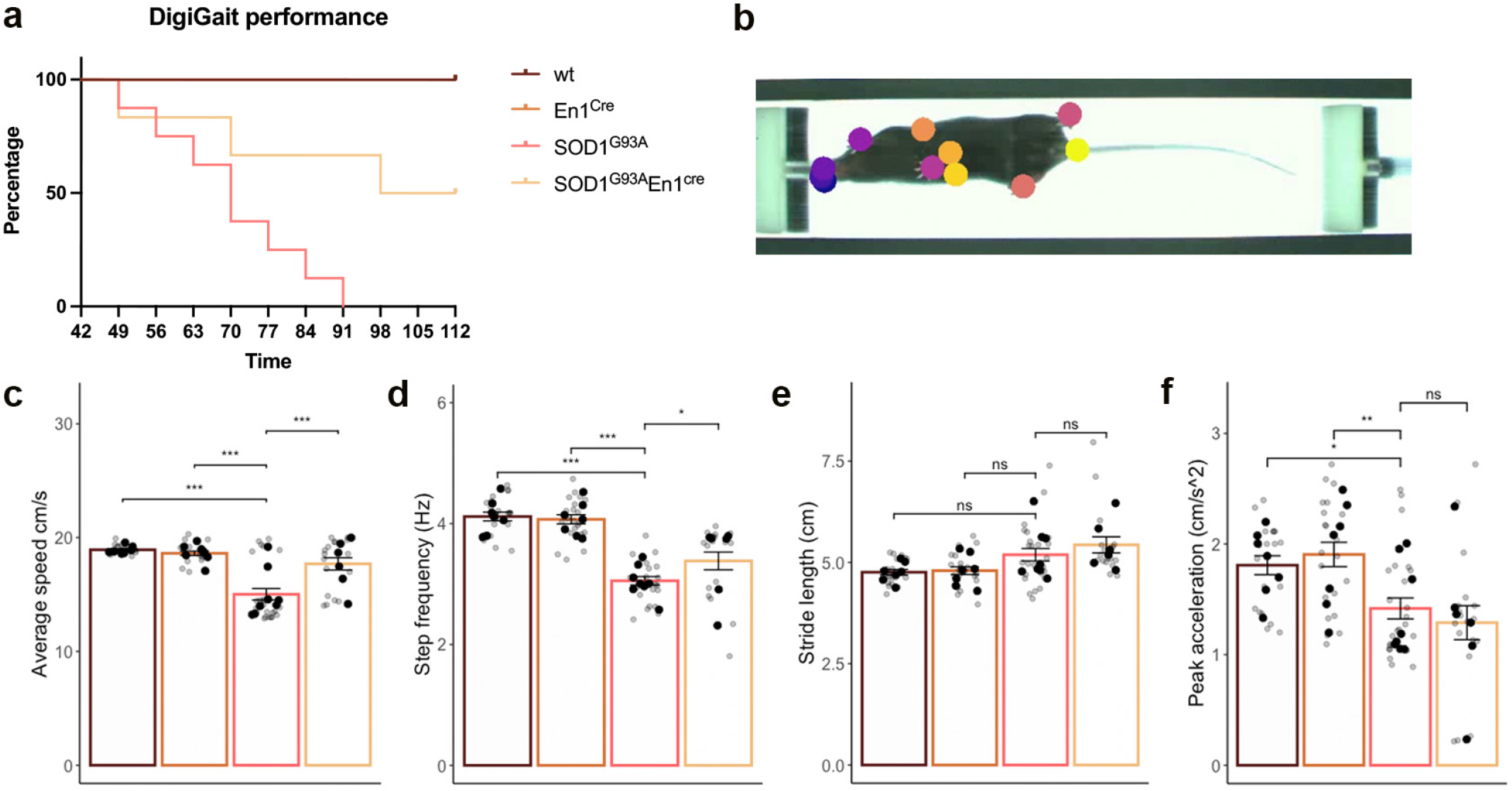
a) Performance of all mice included in the study on a treadmill at a speed of 20 cm/s. Percentage shows that between P49 and P63 37.5% of SOD1^G93A^ mice fail at the task, while only the 16.7% of the SOD1^G93A^;En1^cre^ cannot cope with the speed (SOD1^G93A^ median=70, SOD1^G93A^;En1^cre^ median=105, Log-rank (Mantel-Cox) test df=3, P<0.0001, Chi square=29.02, WT and SOD1^G93A^ N=8 mice, En1^cre^ N=7, SOD1^G93A^;En1^cre^ N=6). b) Example of ventral view tracking of mice on the treadmill, 11 markers were used. c) Differences in average speed (One-way ANOVA and Dunnett’s post hoc, P = 2.5e-05) and d) step frequency (One-way ANOVA and Dunnett’s post hoc, P = 0.033) at onset of locomotor phenotype showing that SOD1^G93A^;En1^cre^ have a later onset when compared to SOD1^G93A^ mice (WT and SOD1^G93A^ N=8 mice, En1^cre^ N=7, SOD1^G93A^;En1^cre^ N=6). Stride length e) and peak acceleration f) remain unchanged at this timepoint. All graphs show mean values ± SEM, averages values in c-f are shown in black, technical triplicates are shown in gray.

**Supplementary figure 3.**
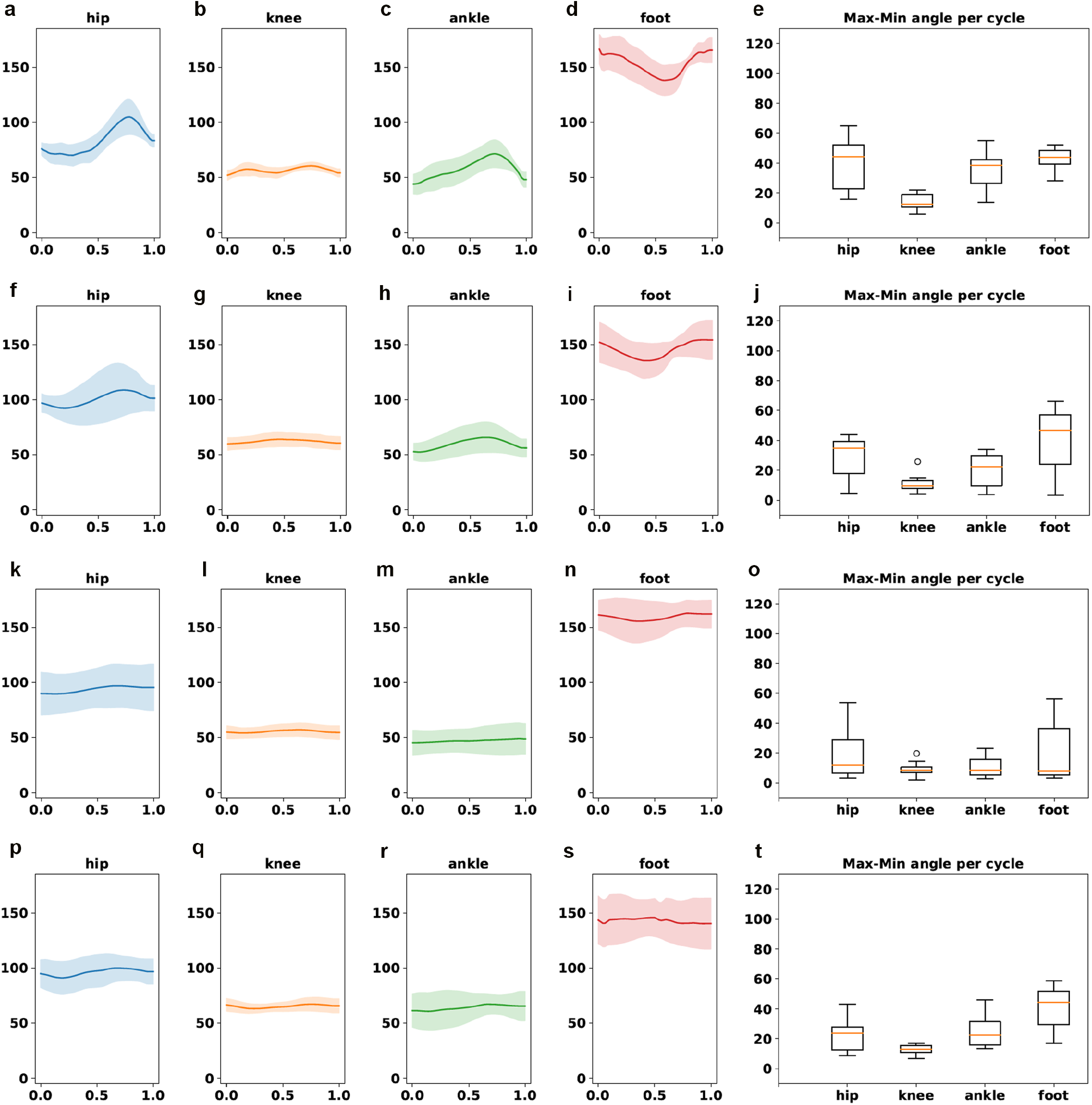
Example of changes in angle amplitudes within one step cycle in WT (a-e), En1^cre^ (f-j), SOD1^G93A^ (k-o) and SOD-1^G93A^;En1^cre^ (p-t) mice. 15 independent steps within each video are utilized for visualization of Max-Min variability (e; j; o; t). Hip k), knee l), ankle m) and foot n) angles are reduced in the SOD1^G93A^ mice at P112 timepoint, indicative of hyperflexion which is exacerbated by disease progression. Changes in amplitude are observed in SOD1^G93A^;En1^cre^ mice overexpressing hEsyt1 (p-t).

**Supplementary video 1.** Kinematic analysis of a SOD1^G93A^ mouse at P112 at a speed of 10 cm/s (average phenotype of SOD1^G93A^ mice at this timepoint). Video was recorded at 150 frames/s, and it is shown at 30 frames/s.

**Supplementary video 2.** Kinematic analysis of a SOD1^G93A^;En1^cre^ mouse at P112 at a speed of 15 cm/s (average phenotype of SOD1^G93A^;En1^cre^ mice at this timepoint), showing differences in hindlimb hyperflexion and in the angle of the mouse body from the belt. Video was recorded at 150 frames/s, and it is shown at 30 frames/s.

**Supplementary table 1.**
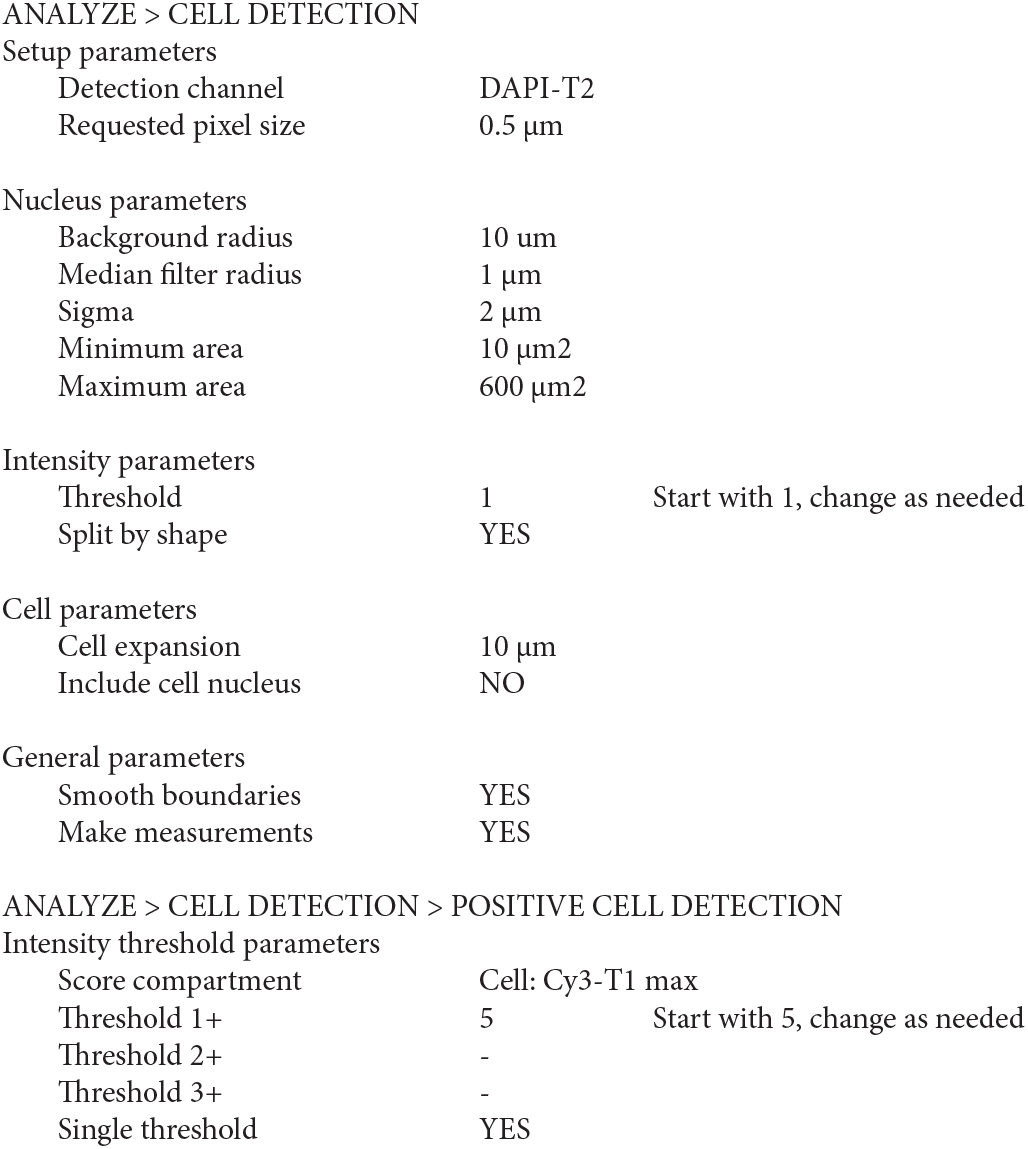
Parameters of QuPath software for RNAscope analysis.

|  |  |
| --- | --- |
| Detection channel | DAPI-T2 |
| Requested pixel size | 0.5 $\mu\text{m}$ |
| Nucleus parameters |  |
| Background radius | 10 $\mu\text{m}$ |
| Median filter radius | 1 $\mu\text{m}$ |
| Sigma | 2 $\mu\text{m}$ |
| Minimum area | 10 $\mu\text{m}^2$ |
| Maximum area | 600 $\mu\text{m}^2$ |

Intensity parameters
|  |  |  |
| --- | --- | --- |
| Threshold | 1 | Start with 1, change as needed |
| Split by shape | YES |  |
| Cell parameters |  |
| Cell expansion | 10 $\mu\text{m}$ |
| Include cell nucleus | NO |
| General parameters |  |
| Smooth boundaries | YES |
| Make measurements | YES |

|  |  |  |
| --- | --- | --- |
| Primary threshold parameters |  |  |
| Score compartment | Cell: Cy3-T1 max |  |
| Threshold 1+ | 5 | Start with 5, change as needed |
| Threshold 2+ | - |  |
| Threshold 3+ | - |  |
| Single threshold | YES |  |

